# Quantification of volumetric morphometry and optical property in the cortex of human cerebellum at micrometer resolution

**DOI:** 10.1101/2021.04.27.441546

**Authors:** Chao J. Liu, William Ammon, Viviana Siless, Morgan Fogarty, Ruopeng Wang, Alessia Atzeni, Iman Aganj, Juan Eugenio Iglesias, Lilla Zöllei, Bruce Fischl, Jeremy D. Schmahmann, Hui Wang

## Abstract

The surface of the human cerebellar cortex is much more tightly folded than the cerebral cortex. Volumetric analysis of cerebellar morphometry in magnetic resonance imaging studies suffers from insufficient resolution, and therefore has had limited impact on disease assessment. Automatic serial polarization-sensitive optical coherence tomography (as-PSOCT) is an emerging technique that offers the advantages of microscopic resolution and volumetric reconstruction of large-scale samples. In this study, we reconstructed multiple cubic centimeters of *ex vivo* human cerebellum tissue using as-PSOCT. The morphometric and optical properties of the cerebellar cortex across five subjects were quantified. While the molecular and granular layers exhibited similar mean thickness in the five subjects, the thickness varied greatly between the crown of the folium and the depth of the fissure in the granular layer within subjects. Layer-specific optical property remained homogenous within individual subjects but showed higher cross-subject variability than layer thickness. High-resolution volumetric morphometry and optical property maps of human cerebellar cortex revealed by as-PSOCT have great potential to advance our understanding of cerebellar function and diseases.

**Highlights:** We reconstructed cubic centimeters of human cerebellar samples at micrometer resolution in five subjects.

Thickness of the granular layer varies greatly between the crowns and depths of cerebellar fissures.

Cross-subject variability is higher in optical property than cortical morphology.

Our results suggest homogenous cell and myelin density in the cortical layers of human cerebellum despite the highly convoluted folding patterns.

## 1. Introduction

The cerebellum contains between 70 and 80% of the neurons in the human nervous system, the vast majority of which are granule cells in the cerebellar cortex (Herculano-Houzel, 2010). The cerebellum is interconnected through afferent and efferent pathways with the cerebral cortex, pontine and olivary precerebellar nuclei, thalamus, and spinal cord. The cerebellum has long been known to be critical for the coordination of movement and it has become apparent from neuroanatomical, electrophysiological, neuroimaging, and clinical studies that its impact on the nervous system is equally important for cognitive and emotional functions (Koziol et al., 2014; Lawrenson et al., 2018; Schmahmann et al., 2019). For example, lesions in the cognitive and limbic regions of cerebellar posterior lobe lead to dysmetria of thought, with impairments of the cerebellar modulation of intellect and emotion manifesting as the cerebellar cognitive affective syndrome (Schmahmann et al., 2019; Schmahmann and Sherman, 1998; Stoodley et al., 2016). There have been fundamental advances in understanding cerebellar corticonuclear connections, circuitry and physiology in animal models (e.g., Apps et al., 2018; De Zeeuw and Ten Brinke, 2015; Voogd and Ruigrok, 2012), but there are still considerable knowledge gaps in understanding cerebellar cortical organization and function in the healthy human brain and how these contribute to pathophysiology and clinical phenomenology in diseases of the cerebellum.

Current volumetric analysis of cerebellar morphometry comes mostly from magnetic resonance imaging (MRI) (Boillat et al., 2018; Sereno et al., 2020). The tightly convoluted pattern of the cerebellar cortex, the limited resolution of MRI, and partial volume effects have impeded comprehensive visualization of the cerebellar folia (Sereno et al., 2020). Optical imaging methods provide superior resolution compared to MRI techniques. Among them, polarization sensitive optical coherence tomography (PSOCT) provides label-free and depth-resolved contrasts that originate from light scattering and tissue birefringence (de Boer et al., 1997). When applied to brain imaging, the reflectivity contrast from PSOCT reveals the gross anatomy and provides quantification of the scattering coefficient of different anatomical structures (Wang et al., 2017; Yang et al., 2020). White matter tracts at microscopic scale can be quantified by polarization-based retardance contrast due to birefringence, an optical property resulting from structural anisotropy (Wang et al., 2011). In particular, the different cortical layers of the cerebellum have showed distinctive optical properties, such as light scattering (Brezinski et al., 1997) and birefringence (Liu et al., 2017). We previously reported automatic serial PSOCT (as-PSOCT) integrated with a tissue slicer and a PSOCT system, which proved to be an effective way to map cubic centimeter samples of human brain (Wang et al., 2018). The as-PSOCT technique makes it possible to examine large-scale cerebellum samples at microscopic resolution.

In this work, we used as-PSOCT to reconstruct cubic centimeters of *ex vivo* human cerebellum samples across five subjects at micrometer scale resolution. The layer thickness and scattering coefficient were quantified to evaluate the morphometric and optical properties of the cerebellar cortex. We further investigated the within-subject and cross-subject variability of cortical layer thickness and scattering coefficient. This quantification may facilitate development of novel biomarkers for neurodegenerative disorders in the cerebellum.

## 2. Material and methods

### 2.1 Human cerebellum samples

Five human cerebellum samples (age at death: 62±10 years, mean ± standard deviation (s.t.d.), 4 males and 1 female) were imaged with as-PSOCT. The brains were obtained from the Department of Neuropathology of the Massachusetts General Hospital (MGH), and all subjects were neurologically normal prior to death. The samples were fixed by immersion in 10% formalin for at least two months. The post-mortem interval did not exceed 24 hours. The *ex vivo* imaging procedures are approved by the Institutional Review Board of the MGH.

### 2.2 System and data acquisition

The as-PSOCT system has previously been described in detail (Wang et al., 2018). The system was composed of an in-house built polarization-maintaining fiber based spectral-domain PSOCT centered at 1300 nm, motorized xyz translational stages, and a vibratome to section the tissue block (Fig. S1). The customized software, written in C++, coordinated data acquisition, xyz stage translation, and vibratome slicing for automatic imaging of tissue blocks. One volumetric acquisition composed of 420 A-lines by 420 B-lines covering a field of view (FOV) of 2.8 mm × 2.8 mm took 3.5 seconds at an A-line rate of 51 kHz. The axial resolution was estimated to be 3.5 μm in tissue. We obtained a lateral resolution of 5.9 μm with a 4 × telescope and a scan lens (LSM03, Thorlabs, Newton, NJ).

The software performed data reconstruction in real time. Inverse Fourier transform of interference-related spectral oscillations yielded complex depth profiles in the form of *A*_1,2_(*z*) exp[*iϕ*_1,2_(*z*)], where *A* and *ϕ* denote the amplitude and phase as a function of depth z, and the subscripts represent the polarization channels. Reflectivity, *R*(*z*), and phase retardance, *δ*(*z*), of the depth profile (A-line), were obtained by *R*(*z*) ∝ *A*_1_(*z*)^2^ + *A*_2_(*z*)^2^, and *δ*(*z*) = arctan[*A*_1_(*z*)/*A*_2_(*z*)], respectively. To increase data acquisition efficiency for large-scale samples, the software saved three-dimensional (3D) reflectivity within a desired depth range for offline scattering coefficient calculations, as well as two-dimensional (2D) *en-face* images of average intensity projection of reflectivity, maximum intensity projection of reflectivity, average retardance, and dominant axis orientation.

Due to the FOV limit in one acquisition, we imaged the block face surface via tile scans with 50% overlapping. We stitched the tiles in the sagittal plane to obtain the volumetric reflectivity data as well as the *en-face* retardance images of each physical slice. A 100 μm thick slice was removed from the tissue surface by the vibratome to expose the deeper region until the whole volume was imaged. The *en-face* images of individual slices were stacked together to render the cubic centimeters of cerebellum tissue.

### 2.3 Cortical layer and white matter segmentation

Segmentation of the cerebellar cortex was performed using a semi-automated method. Manual segmentation was first conducted on one out of five adjacent sagittal slices in the Freeview software (part of the FreeSurfer software package) (Fischl, 2012). The boundaries of the molecular layer, the granular layers, and the white matter were drawn based on the volumetric retardance images, and a label was created for each of the structure. An automated algorithm called SmartInterpol was then applied to complete the rest of the slices (Atzeni et al., 2018). SmartInterpol uses a combination of label fusion and deep learning methods to transfer the segmentations from the labeled to the unlabeled slices, and has two major advantages: (i) reducing the burden of manual delineation; and (ii) producing segmentations that are smooth in the orthogonal views, which is very difficult to achieve with completely manual tracing. The label fusion component of SmartInterpol relies on affine and nonrigid registration methods available in the publicly available NiftyReg package (Modat et al., 2010; Ourselin et al., 2001), whereas the deep learning component relies on a fully convolutional network (Shelhamer et al., 2016). The segmentation results were used to compute the layer thickness for the cerebellum as described in the following section.

### 2.4 Quantification of morphometric and optical properties

#### Layer thickness estimation

A voxel-based algorithm was applied to estimate the thickness of the molecular and granular layers of the cerebellar cortex (Aganj et al., 2009). For every voxel in the volumetric data, the algorithm considered lines passing through the voxel inside the segmented layer (in 193 3D directions orientations uniformly distributed on the sphere), computed the length of each line intersecting with the layer boundaries, and selected the shortest length as the estimated thickness. To facilitate further analysis of layer thickness, we used the thickness of the skeleton of the segmented structure. The skeleton of the cortex was defined as the two-voxel-wide mask positioned in the middle of the layer, far from the inner/outer surfaces (Aganj et al., 2009).

#### Optical properties

We quantified the voxel-wise attenuation coefficients following the method of Vermeer et al. In the near-infrared spectral range, light attenuation within the tissue is dominated by scattering, whereas absorption is negligible. Therefore, we used the scattering coefficient (μ_*s*_), calculated as 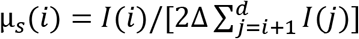 to represent the optical properties, where *i* is the pixel number in depth, *I* is the reflectivity signal, Δ is the pixel size in depth, and *d* is the total imaging depth (Vermeer et al., 2013).

#### Variation index

Variation index was a term we defined in the study, which represented the heterogeneity of layer thickness or μ_*s*_ within a subject. It was calculated as *Variation Index* = 〈*Q*_*upper*,15%_(*X*)〉/〈*Q*_*lower*,15%_(*X*)〉, where *X* represents the measurement (layer thickness or μ_*s*_), 〈 〉 is the mean operation, *Q*_*upper*,15%_ is the top 15% of the measurements and *Q*_*lower*,15%_ is the bottom 15% of the measurements. A variation index ≈ 1 indicates a uniform distribution in the measurement, whereas a variation index ≫ 1 indicates high heterogeneity. The variation index of layer thickness reflects the extent of convolution of the cerebellar cortex and is equivalent to the folding index of a specific layer.

### 2.5 Correlation between morphometric and optical properties

We calculated correlation coefficients between the layer thickness and μ_*s*_ in the molecular and granular layers, to evaluate the correlation between morphometric and optical properties. The greater number of voxels in the folia compared to the fissures, specifically in the granular layer, potentially leaned towards a folial bias; and therefore, we used the layer thickness and μ_*s*_ on the skeleton for the correlation analysis.

## 3. Results

### 3.1 Reconstruction of human cerebellum samples at micrometer-scale resolution

Each of the five human cerebellum samples imaged by as-PSOCT was about 4 cm^3^. The sample shown in Fig. 1 was 2.8 × 1.8 × 0.75 cm^3^. We first reconstructed the *en-face* retardance images of the 75 slices and stacked them together to recover the whole cerebellar volume (Fig. 1A). The white matter showed higher retardance values compared with the cerebellar cortex. Furthermore, the granular layer showed higher retardance than the molecular layer, in agreement with the findings in previous studies (Liu et al., 2017; Wang et al., 2018). We then calculated the μ_*s*_ map of each slice by averaging depth-wise μ_*s*_ and stacked them together to represent the volumetric reconstruction. The coronal, axial, and sagittal planes of the μ_*s*_ images revealed similar anatomical structure to the retardance images (Fig. 1B). The molecular layer, granular layer, and white matter showed distinctive μ_*s*_ characteristics.

**Fig. 1.**
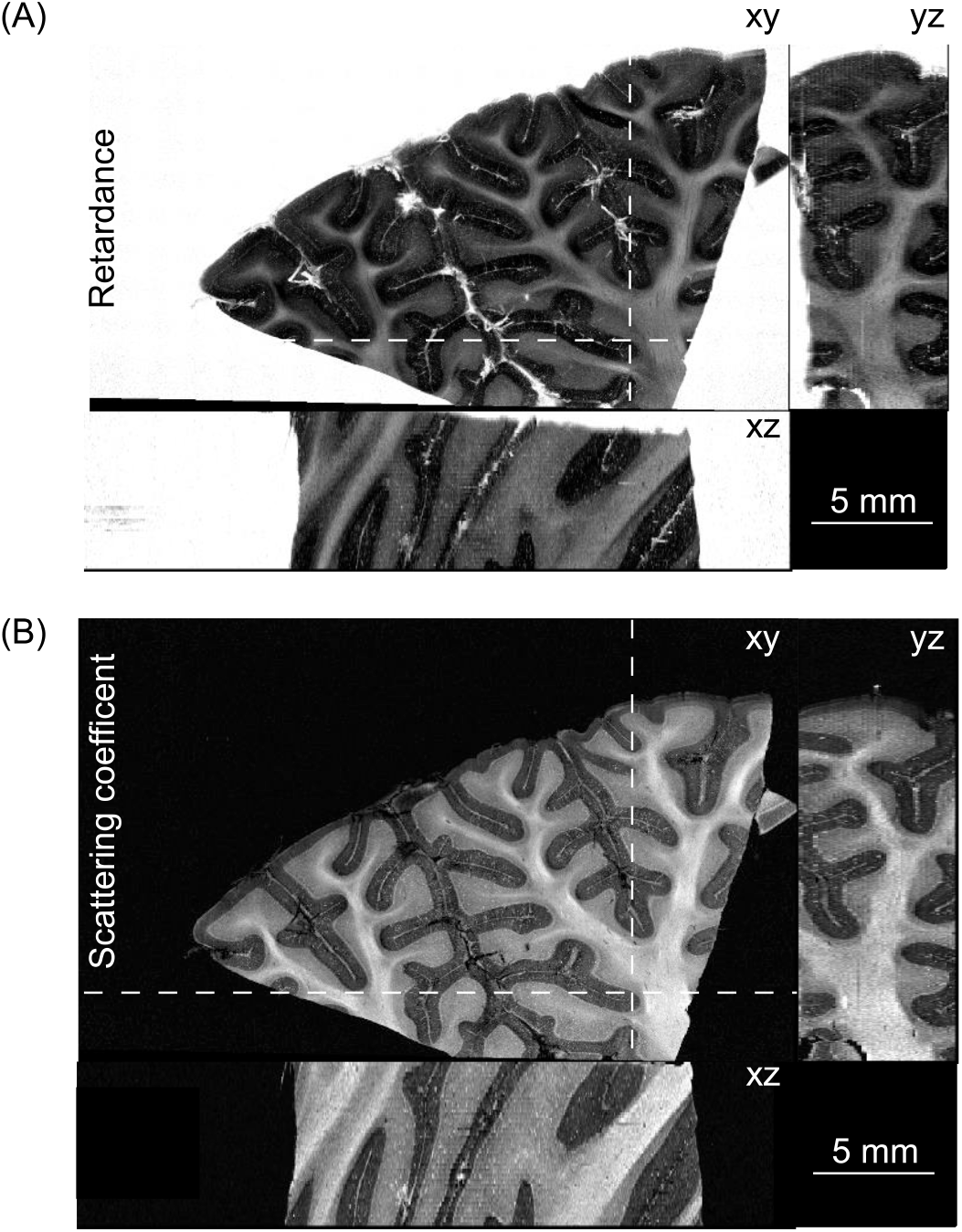
Volumetric reconstruction of human cerebellar tissue (2.8 × 1.8 × 0.75 cm^3^) from as-PSOCT. The orthogonal views of (A) retardance images and (B) scattering coefficient maps are shown. The locations of the axial (xz) and coronal (yz) planes in A and B are indicated by the dashed lines on xy-plane. Scale bars: 5 mm.

The prominent retardance contrast allows segmentation of the cerebellar cortex and the white matter, with the former further divided by cortical laminar structures including the molecular and granular layers (Fig. 2). Due to the resolution limit of the down-sampled reconstruction, we did not segment the interleaved Purkinje cell monolayer. Remarkably, as mentioned in Section 2.3 above, the semi-automated segmentation reduced the labor time by a factor of five and also yielded less inter-slice variability at structural boundaries where subtle contrast challenged the consistency of human judgement in the full manual segmentation (Fig. 3). As a result, the volumetric segmentations were inherently smooth. The structural segmentation enables further analysis of morphometric and optical properties in a layer-specific manner.

**Fig. 2.**
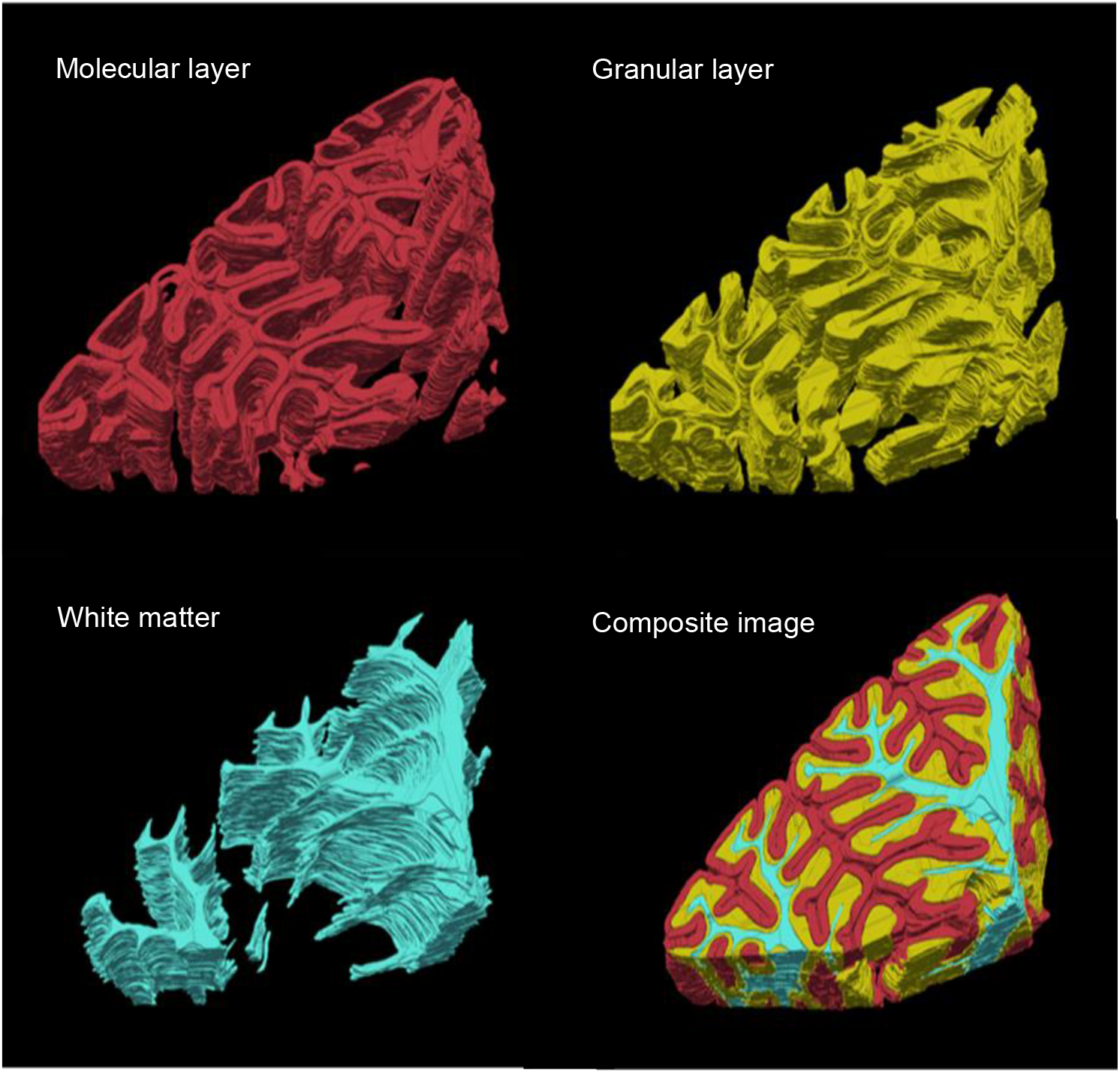
Volume rendering of segmented molecular layer (red), granular layer (yellow) and white matter (cyan) for the cerebellar lobules. The whole segmented volume is shown in the composite image of the three structures.

**Fig. 3.**
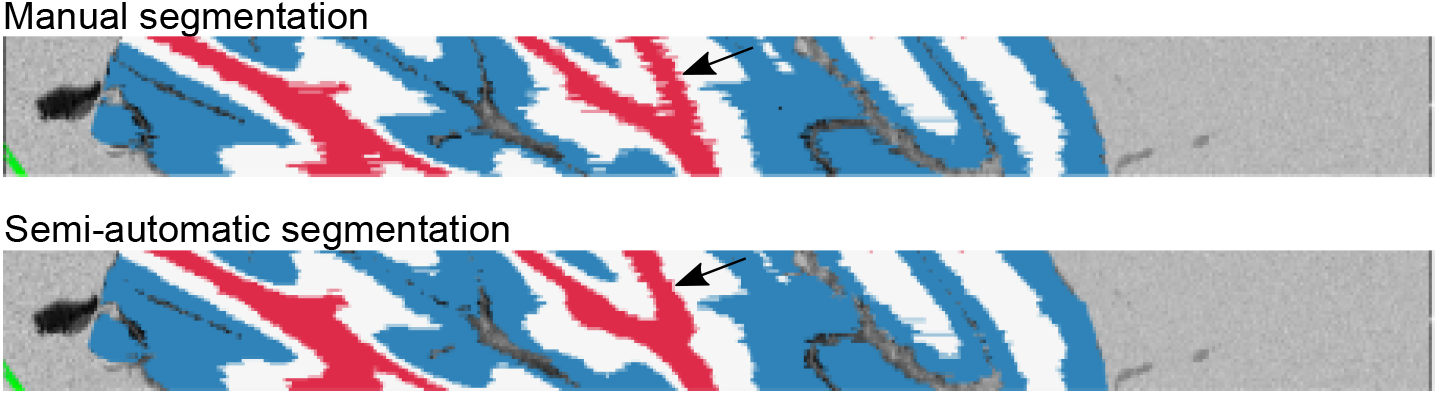
Comparison of full manual (top) and semi-automatic (bottom) cerebellum segmentation on a cross-slice viewing plane. Black arrows highlight the inconsistency from manual segmentation and the smoothness from semi-automatic segmentation.

### 3.2 Morphometric and optical properties of human cerebellar cortex

We quantified the thickness of the molecular and granular layers from the five cerebellar samples. The thickness maps of the molecular and granular layers were obtained using a voxel-based algorithm and visualized in one cerebellar volume as an example (Fig. 4A). The value at every voxel represents the thickness of the corresponding layer calculated at that voxel. We observed that the granular layer showed much greater thickness in the crown of the folium compared to the values in the depth of the fissure, due to the convoluted patterns of the cerebellar cortex (see enlarged box in Fig. 4A). In contrast, the thickness of the molecular layer was more uniform.

**Fig. 4.**
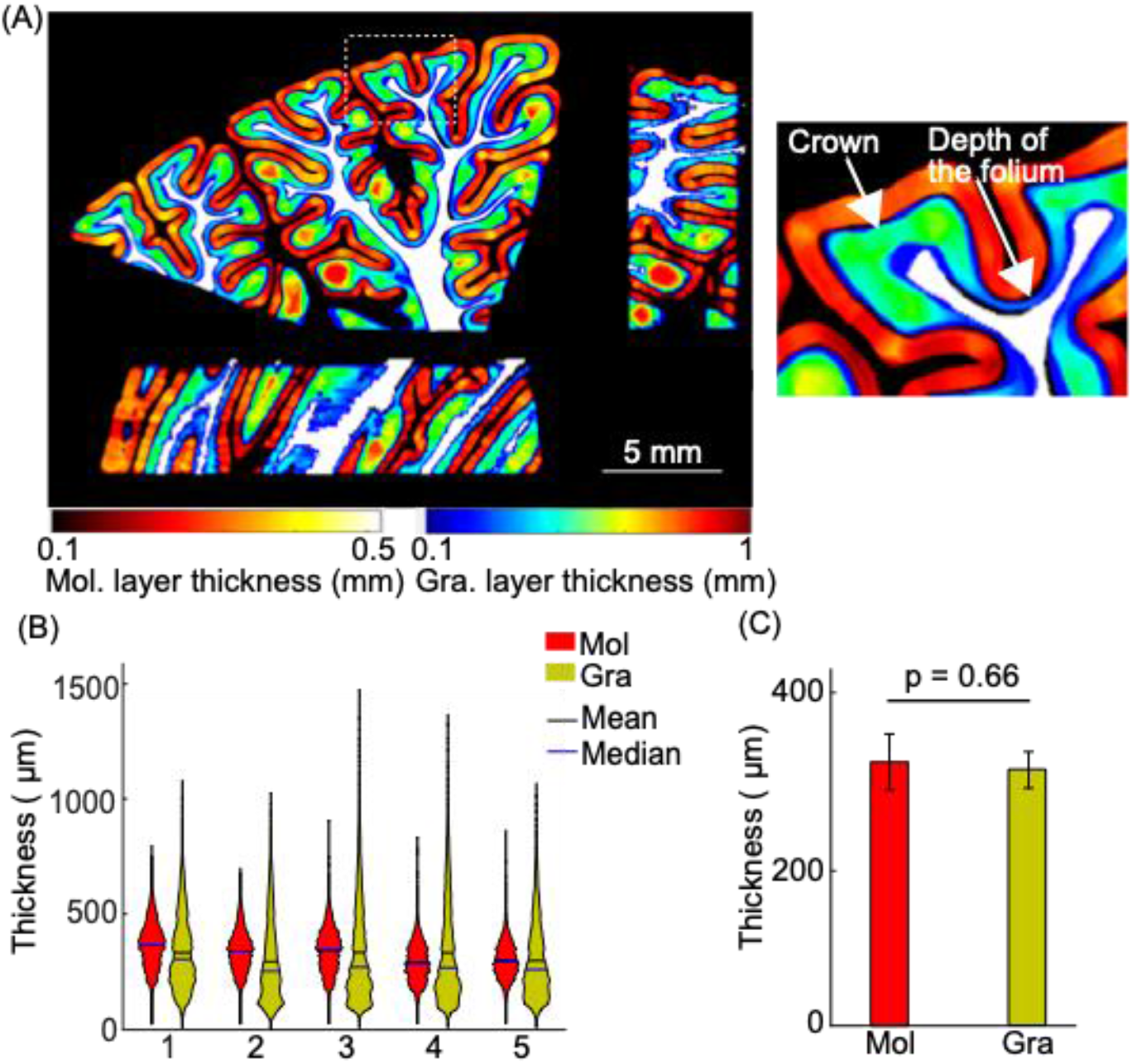
Cortical layer thickness in five cerebellar samples. (A) Molecular and granular layer thickness estimation in orthogonal views. The molecular and granular layer thicknesses are indicated by the hot and jet color bars, respectively. White matter is shown in white for anatomical reference. The enlarged box on the right panel highlights the granular layer in the crown and depth of the folium (white arrows). Scale bar: 5 mm. Violin plots of molecular and granular layer thickness in individual subjects. The width and height of the violin plots correspond with the frequency and the value of thickness measurements. The mean and median of the measurements are also shown in the plots as black and blue solid lines. (C) Average cortical layer thickness of the five subjects. Error bars indicate standard deviations. Mol: molecular layer; Gra: granular layer.

We examined the thickness distribution of the molecular and granular layers in the five subjects (Fig. 4B). The distribution within a layer was consistent across all subjects, while the distributions of the thickness in the molecular and granular layers revealed distinct patterns. The distribution of thickness of the granular layer was skewed to the lower end while having a long tail towards higher thickness, whereas the distribution of thickness of the molecular layer was narrower and more symmetric. The molecular and granular layers shared similar mean thickness, 322 ± 32 μm and 319 ± 20 μm respectively (Fig. 4C, mean ± s.t.d., p = 0.66, paired-sample *t*-test, n = 5 subjects).

Tissue microstructure changes the light scattering events and thus manifests in the optical properties. After obtaining the scattering coefficient map (Fig. 1B) and the volumetric segmentation (Fig. 2), we further analyzed μ_*s*_ in the molecular layer, the granular layer, and the interior of the white matter. Within individual subjects, white matter had the highest μ_*s*_, followed by the granular layer and the molecular layer (Fig. 5A). We then evaluated μ_*s*_ across five subjects and found consistent patterns, with μ_*s*_ of 3.3 ± 0.5 mm^−1^, 4.6 ± 0.6 mm^−1^, and 5.7 ± 1.3 mm^−1^ in molecular layer, granular layer, and white matter, respectively (Fig. 5B, mean ± s.t.d., p = 0.0041, one-way ANOVA, n = 5 subjects). Due to the effect of speckle noise in OCT imaging, we observed extremely high and low values in the μ_*s*_ data (Fig. S2A). We excluded the μ_*s*_ values outside the three-sigma confidence intervals for the further analysis; however, the exclusion did not change the μ_*s*_ pattern in these structures and the statistical analysis (Fig. S2B). The folial white matter is composed of bundles of myelinated axons. Consequently, the high scattering in the white matter is associated with optical properties of myelin compared to cortex (Wang et al., 2011). Within the cerebellar cortex, the granular layer is densely packed with granule cells and contains myelinated axons in the adult brain (Liu et al., 2017).

**Fig. 5.**
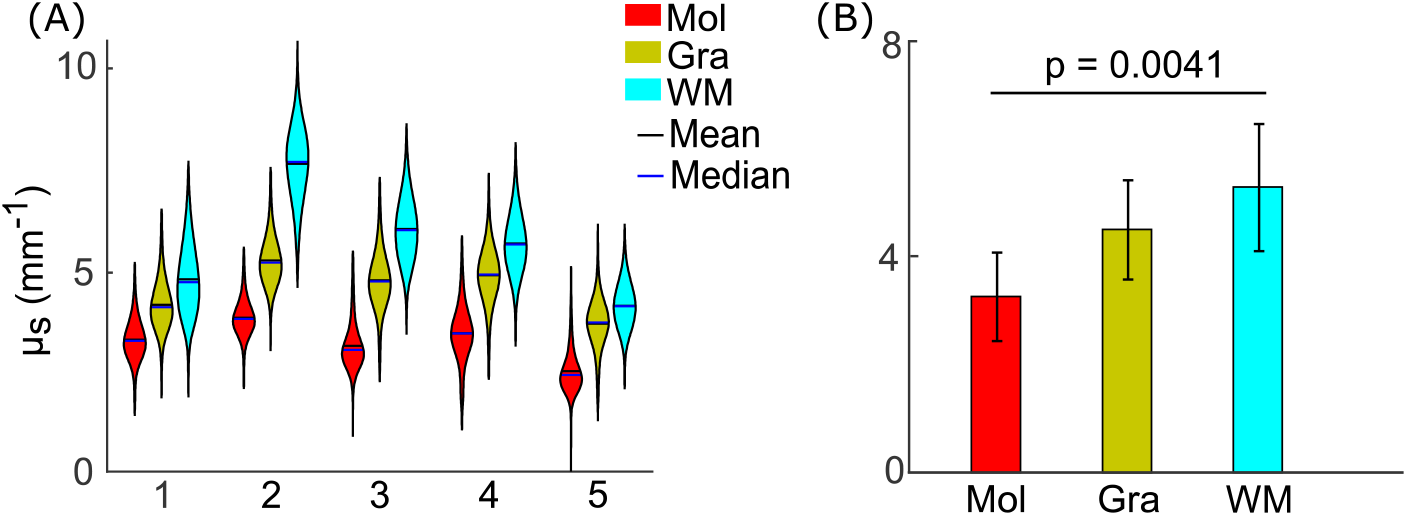
Optical scattering coefficient measurements in five cerebellar samples. (A) Violin plots of μ_*s*_ measurements in molecular layer, granular layer, and white matter in individual subjects. The width and height of the violin plots correspond with the frequency and the value of μ_*s*_ measurements. The mean and median of the measurements are represented as black and blue solid lines. (B) Average μ_*s*_ measurements in molecular layer, granular layer, and white matter across the five subjects. Error bars indicate standard deviations. Mol: molecular layer; Gra: granular layer; WM: white matter.

### 3.3 Within-subject variation of morphometric and optical properties in cerebellar cortex

To further understand the heterogeneity of morphometric and optical properties within individual layers, we evaluated the folding index of morphology and variation index of μ_*s*_ in the cerebellar cortex. Since layer thickness has the highest values at the crown of the folium and the lowest in cortex situated at the depth of the fissure (see Fig. 4A), the folding index was a marker of the extent of cortical folding in human cerebellum, by a ratio of the thickness between the two sets of locations (top 15% and bottom 15% of thickness measurement). We found that granular layer thickness varied by a factor of almost four. In contrast, the variation of molecular layer thickness was lower, close to two. The folding indices between the two layers were significantly different (Fig. 6A, 4.0 ± 0.6 and 1.8 ± 0.07, respectively, mean ± s.t.d., p = 7e−5, paired-sample *t*-test, n = 5 subjects). In contrast to the pronounced variations in morphology, the optical property is more spatially uniform within a layer regardless of crown or trough, with a variation index close to one. The variation indices of μ_*s*_ in the molecular layer and the granular layer were similar (Fig. 6B, 1.3 ± 0.04 and 1.3 ± 0.02, respectively, mean ± s.t.d., p = 0.61, paired-sample *t*-test, n = 5 subjects).

**Fig. 6.**
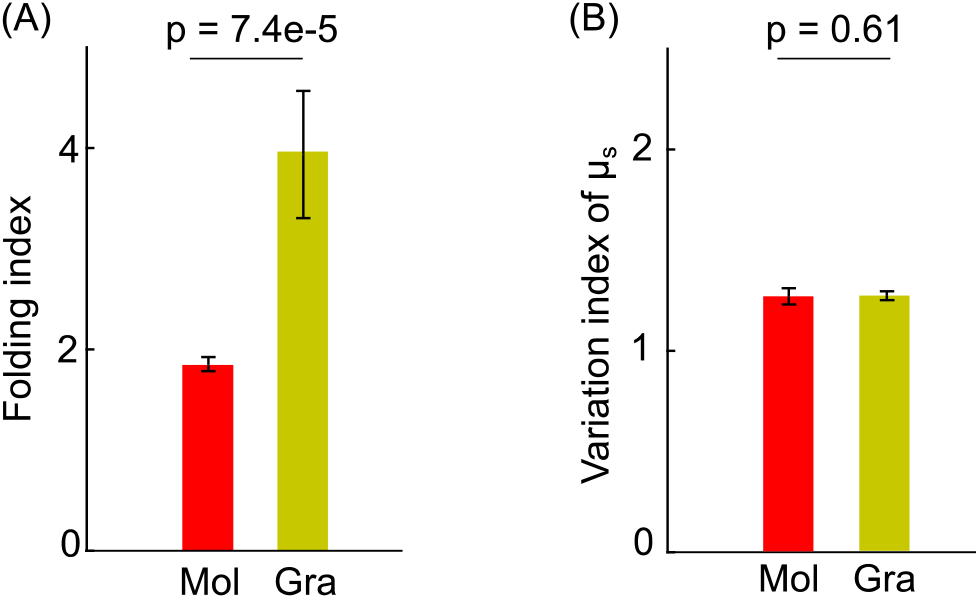
The folding index (A) and the variation index of μ_*s*_ (B) in the molecular and granular layers. Error bars indicate standard deviations among the five subjects. Mol: molecular layer; Gra: granular layer.

### 3.4 Cross-subject variation of morphometric and optical properties in the cerebellar cortex

The results in Section 3.2 show that the mean layer thickness was comparable across the five subjects (Fig. 4C), while the mean optical properties exhibited greater variations (Fig. 5B). To quantitatively evaluate the cross-subject variability, we calculated the coefficient of variation (CV, equal to the standard deviation divided by the mean) from the mean layer thickness and layer-specific mean μ_*s*_ across the five subjects. The CV of the mean layer thickness in the molecular and granular layers was 9.8% and 6.3%, respectively. In contrast, we found the CV of the mean μ_*s*_ to be higher than that of the mean layer thickness, resulting in 15.2% and 13.0% in the molecular and granular layers, respectively. It is noted that despite the highly convoluted patterns, the mean thickness of a layer differed by less than 10% across the subjects, indicating a low cross-subject variability in mean thickness of human cerebellar cortex within the age range of this study.

### 3.5 Correlation between morphometric and optical properties in the cerebellar cortex

We analyzed the correlation between layer thickness and μ_*s*_ by quantifying the correlation coefficients (R) between these two measures. The layer thickness exhibited little correlation with μ_*s*_ in either the molecular or granular layers of the five cerebellar samples (Fig. 7). The correlation coefficient with the largest magnitude that we observed was 0.1. The contours in Fig. 7 illustrate the 2D distribution of thickness – μ_*s*_ with correlation coefficients shown for every subject. Overall, no significant correlation was found between layer thickness and optical properties.

**Fig. 7.**
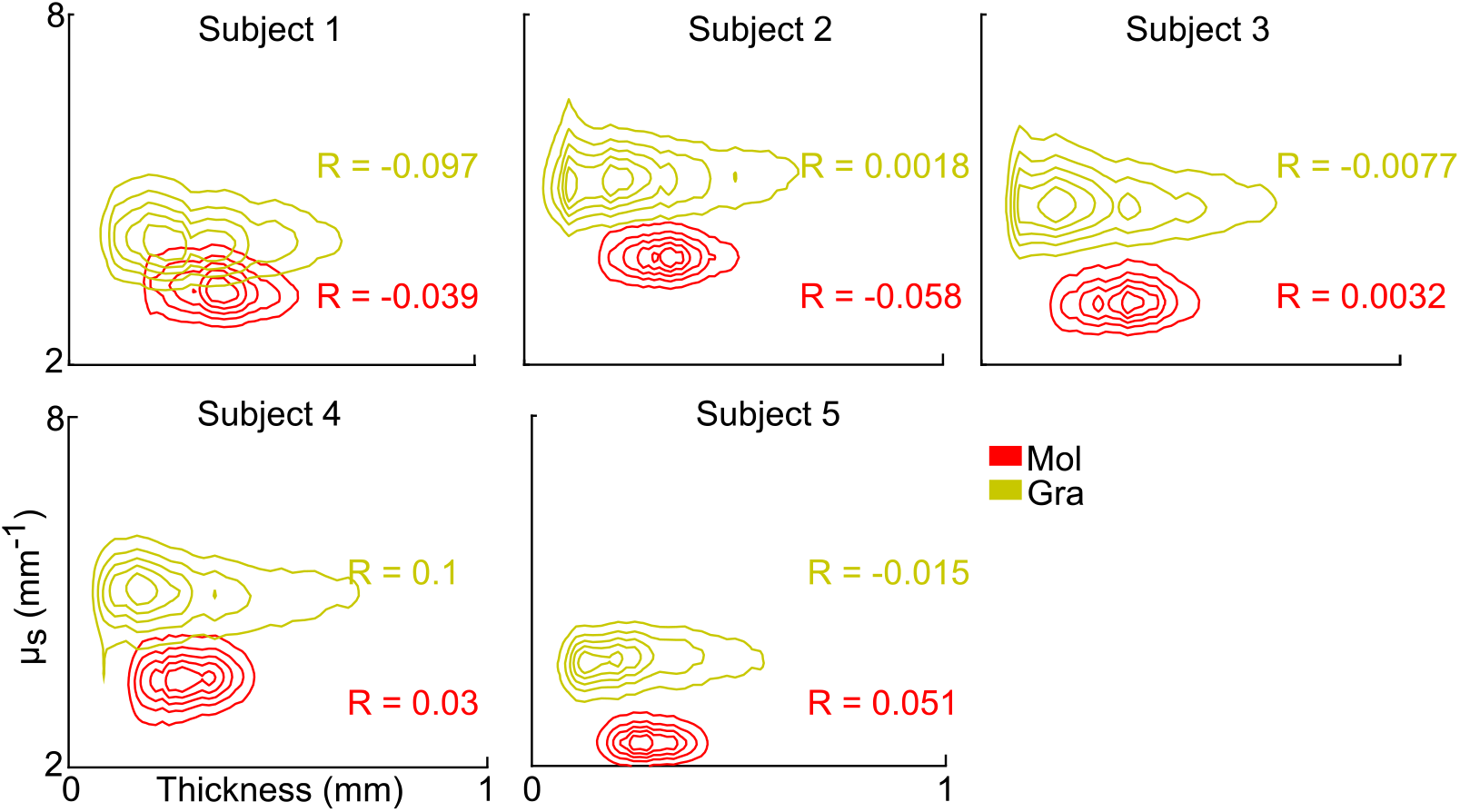
Contour plots of 2D distribution between thickness and μ_*s*_ measurements in the molecular and granular layers of five cerebellar samples. The correlation coefficients (R) between layer thickness and μ_*s*_ are shown for each subject in molecular and granular layers. Mol: molecular layer; Gra: granular layer.

## 4. Discussion

Over 100 years after Santiago Ramón y Cajal’s pioneering work in identifying neurons and axonal tracts, human brain mapping at micrometer resolution still remains challenging. The next generation of optical imaging methods requires rapid acquisition rates and volumetric reconstruction in large-sized sample. The as-PSOCT technique supports 3D reconstruction of cubic centimeters of human brain samples at 3-10 μm resolution. In this study, we presented the quantification of morphometric and optical properties in large cerebellum volumes using as-PSOCT.

To characterize the morphometry of the human cerebellar cortex, we first measured cortical layer thickness from the volumetric reconstruction (Fig. 4). MRI techniques have been used to estimate total cerebellar volume (Hara et al., 2016) and visualize the cerebellar cortical layers in-plane (Marques et al., 2010). However, the resolution of MRI limits its ability to measure the layer thickness in the cerebellum. By using as-PSOCT, we showed that the mean thickness of both the molecular layer and the granular layer is ~320 μm, and the thickness in the depth of the fissure was smaller than one pixel even with state-of-the-art 100 μm resolution *ex vivo* MRI technique (Edlow et al., 2019). Quantification of the folding morphology is important in order to establish biomarkers from neuroimaging data, especially in neurological diseases (Mangin et al., 2010). Current gyrification measurements from MRI (Schaer et al., 2012) have been applied only to the cerebral cortex, while the small folia in the cerebellar cortex have been left unexplored. In this study, we showed that with high resolution, as-PSOCT provides a viable solution to measuring the folding index of cerebellar cortex in a layer-specific manner (Fig. 6A). We found a high folding index in the granular layer that had not been reported before. The high-quality data generated by as-PSOCT can be used to fit MR models to construct a neuroarchitectural map of the human cerebellum, an approach not available using current imaging techniques.

Our as-PSOCT system allows quantification of a scattering coefficient map. We found different scattering characteristics in the laminar structures (Figs. 1B and 5). Further analysis revealed significantly different scattering coefficient between the molecular layer and the white matter across the five subjects (p = 0.0036, one-way ANOVA with Dunn-Sidak correction, n = 5 subjects); however, we did not find a significant difference between the molecular and granular layers (p = 0.11, one-way ANOVA with Dunn-Sidak correction, n = 5 subjects), possibly related to a limited sample size. The optical properties generated by as-PSOCT enabled the quantitative assessment of microstructural features of cerebellar tissues, which may serve as a potentially useful tool for the evaluation of pathological process. Many cerebellar neurodegenerative diseases include loss of Purkinje neurons and atrophy of the cerebellar cortex, such as multiple system atrophy and the spinocerebellar ataxias (Koeppen, 2018). Optical scattering changes have been observed in neurodegenerative disease (Hanlon et al., 2008), especially in spinocerebellar ataxia type 1 (Liu et al., 2018). Therefore, the scattering coefficient may serve as a new biomarker of the neuropathology in cerebellar neurodegeneration, including alterations in myelin cell and density.

In the samples from the five subjects we found the CV of the mean μ_*s*_ to be 15.2% and 13.0% in the molecular and granular layers, respectively, which were higher than the CV of mean layer thickness (<10%). Therefore, we observed higher cross-subject variability in the optical properties than in the morphology for our five samples. These results suggest that optical properties may be a sensitive marker of neuropathology, identifiable before large-scale histopathological changes can be detected.

The homogeneous optical properties within the molecular and granular layers and the lack of correlation with thickness of the layers is a novel observation (Figs. 6B and 7), suggesting a uniformity of cellular and myelin density despite the pronounced folding patterns in the cerebellar cortex (Fig. 6A). Gyral and sulcal morphological analysis reported by MRI has focused on the impact of neurological diseases on the cerebrum (Graham and Sharp, 2019; Im et al., 2008). In a topographical analysis of demyelination and neurodegeneration in multiple sclerosis subjects, for example, Haider et al. found heterogeneous neurodegeneration betweeb the crowns and the sulci (Haider et al., 2016). With as-PSOCT, similar analyses can be performed in the cerebellum to explore a variety of neurodegenerative diseases. For example, it is possible that the cortex at the crowns and depths of the cerebellar folia are affected differently in spinocerebellar ataxia type 1, in which the Purkinje cell layer and granular layer show heterogeneous characteristics of neuronal loss (Genis et al., 1995). The analysis of morphometric and optical properties of the human cerebellum opens new avenues to investigate spatial-structural patterns of neuropathology in cerebellar disorders.

In the current study we used a low numerical aperture objective to image the cerebellar samples. Due to the limit of lateral resolution, we did not analyze the thin Purkinje cell layer. However, it is important to acknowledge that Purkinje cells are a critical information processing hub in the cerebellum. The as-PSOCT system has the potential to incorporate high numerical aperture objectives to visualize Purkinje cells and other cells (Magnain et al., 2015; Wang et al., 2016). Further developments of our technique are under way to provide this level of cellular information necessary for the study of neurological diseases.

## 5. Conclusions

We used as-PSOCT to reconstruct volumetric representations of the human cerebellum and quantify morphometric and optical properties in post-mortem cerebellar tissue in five samples. We recovered the laminar structure of the folia, which cannot be achieved by MRI. The laminar structures in the cerebellar cortex were differentiated by both retardance and scattering coefficient contrasts. The layer thickness and folding index revealed a complex folding pattern in the cerebellar cortex, where the thickness varied greatly in the crown of the folium compared to that in the depth of the fissure in the granular layer. The thickness in the molecular layer was less variant. We also found a uniform distribution of cell and myelin content reflected in the homogeneous optical properties within a cortical layer despite the complicated folding patterns. Conversely, the higher cross-subject variability for optical properties than for cortical thickness in the cerebellum suggested that the optical properties may serve as a more sensitive parameter to evaluate pathological conditions. High-resolution as-PSOCT data supports a detailed atlas to be built in the cerebellum that will enable *in vivo* prediction in MRI assessment that would not otherwise be possible. Quantification in morphometric and optical properties may serve as new biomarkers in cerebellar diseases such as multiple system atrophy and spinocerebellar ataxias, and therefore have the potential to influence pathological evaluations in scalable population size.

## Data and code availability statement

Manual segmentation tool of Freeview is available as part of the open-source software FreeSurfer: https://surfer.nmr.mgh.harvard.edu/fswiki/DownloadAndInstall. The layer thickness code is available at https://www.nitrc.org/projects/thickness. Data used in this work are available upon request.

## Acknowledgements

We greatly appreciated the insightful discussions with Dr. Douglas Greve for this work. This work was supported by the National Institutes of Health (NIH), specifically: in part by the BRAIN Initiative Cell Census Network grant (U01MH117023), the National Institute for Biomedical Imaging and Bioengineering (R00EB023993, R21EB018907, U01EB026996, R01EB019956, R01EB006758, R01EB023281, P41EB030006, P41EB015896), the National Institute of Mental Health (R01MH123195, R01MH121885, RF1MH123195), the National Institute on Aging (R01AG064027, R01AG008122, R01AG016495, R01AG070988, R56AG068261, R56AG064027), the National Institute of Diabetes and Digestive and Kidney Diseases(K01DK101631), and the National Institute for Neurological Disorders and Stroke (R01NS0525851, R21NS072652, R01NS070963, R01NS083534, 5U01NS086625, 5U24NS10059103, R01NS105820), as well as Shared Instrumentation Grants (S10RR023401, S10RR019307, and S10RR023043). Additional support was provided by the NIH Blueprint for Neuroscience Research (5U01MH093765), part of the multi-institutional Human Connectome Project, as well as by the European Research Council (Starting Grant 677697, project “BUNGEE-TOOLS”) and by Alzheimer’s Research UK (Interdisciplinary Grant ARUK-IRG2019A-003).

