## Supplementary material for "Quantification of volumetric morphometry and optical property in the cortex of human cerebellum at micrometer resolution": Fig. S1, Fig. S2

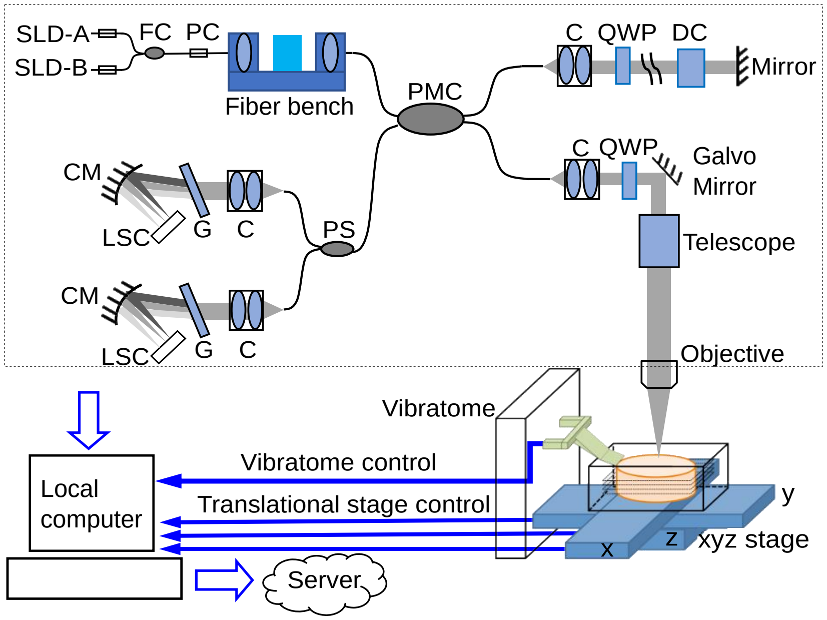


Fig. S1 Schematic of as-PSOCT system. The dashed box indicates the homemade spectral domain PSOCT. Abbreviations: SLD, super luminescent diode; FC, fiber coupler; PC, polarization controller; PMC, polarization-maintaining fiber coupler; C, collimator; QWP, quarter-wave plate; DC, dispersion compensation block; PS, polarization splitter; CM, concave mirror; G, grating; LSC, line scan camera.

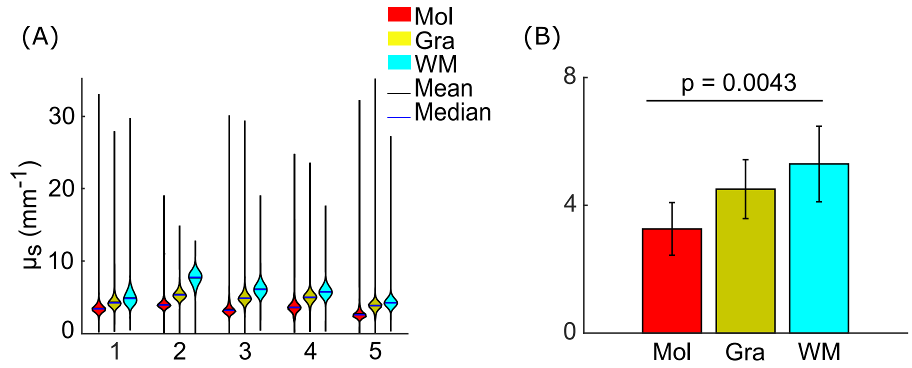


Fig. S2 Optical scattering coefficients in five cerebellum samples without data exclusion. (A) Violin plots of $\mu_s$ measurements in molecular layer, granular layer, and white matter in individual subjects. The width and height of the violin plots correspond with the frequency and the value of $\mu_s$ measurements. The mean and median of the measurements are represented as black and blue solid lines. (B) Average $\mu_s$ measurements in molecular layer, granular layer, and white matter across the five subjects. Error bars indicate standard deviations. Mol: molecular layer; Gra: granular layer; WM: white matter.
